# Preliminary Immunogenicity of a Pan-COVID-19 T Cell Vaccine in HLA-A*02:01 Mice

**DOI:** 10.1101/2021.05.02.442052

**Authors:** Brandon Carter, Jinjin Chen, Clarety Kaseke, Alexander Dimitrakakis, Gaurav D. Gaiha, Qiaobing Xu, David K. Gifford

**Affiliations:** MIT Computer Science and Artificial Intelligence Laboratory, Cambridge, MA, USA; Department of Biomedical Engineering, Tufts University, Medford, MA, USA; Ragon Institute of MGH, MIT and Harvard, Cambridge, MA, USA; MIT Electrical Engineering and Computer Science, Cambridge, MA, USA; MIT Biological Engineering, Cambridge, MA, USA

**Author notes:** These authors contributed equally to this work (Brandon Carter, Jinjin Chen).

## Abstract

New strains of SARS-CoV-2 have emerged, including B.1.351 and P.1, that demonstrate increased transmissibility and the potential of rendering current SARS-CoV-2 vaccines less effective. A concern is that existing SARS-CoV-2 spike subunit vaccines produce neutralizing antibodies to three dimensional spike epitopes that are subject to change during viral drift. Here we provide an initial report on the hypothesis that adaptive T cell based immunity may provide a path for a pan-COVID-19 vaccine that is resilient to viral drift. T cell based adaptive immunity can be based on short peptide sequences selected from the viral proteome that are less subject to drift, and can utilize multiple such epitopes to provide redundancy in the event of drift. We find that SARS-CoV-2 peptides contained in a mRNA-LNP T cell vaccine for SARS-CoV-2 are immunogenic in mice transgenic for the human HLA-A*02:01 gene. We plan to test the efficacy of this vaccine with SARS-CoV-2 B.1.351 challenge trials with HLA-A*02:01 mice.

## 1 Introduction

Concern has been raised by the observations that new strains of SARS-CoV-2 are more transmissible (Davies et al., 2021; Tegally et al., 2021), and observed spike variants tested as pseudoviruses are more resilient to neutralization by sera produced by spike subunit vaccines (Liu et al., 2021b; Wu et al., 2021), and more resilient to neutralization by convalescent sera (Wibmer et al., 2021). The ultimate clinical implications of these findings are at present unclear, but they have motivated a search for vaccination methods that are resilient to new strains of SARS-CoV-2.

We hypothesize that an alternative to vaccine-induced antibody-based viral neutralization is the production of a robust cellular immune response to protect an individual against symptomatic SARS-CoV-2 infection. We have proposed T cell vaccines that are predicted to produce both CD8^+^ and CD4^+^ T cell responses (Liu et al., 2020, 2021a), and here we present the initial immunogenic evaluation of such a vaccine in a HLA-A*02:01 human transgenic mouse model.

## 2 Methods

### 2.1 Vaccine peptide selection

#### MHC Class I

Eight peptides from SARS-CoV-2 were selected as a subset of the MHC class I *de novo* MIRA only vaccine design of Liu et al. (2021a). We filtered this set of 36 peptides to the 8 peptides predicted to be displayed by HLA-A*02:01 by a combined MIRA and machine learning model of peptide-HLA immunogenicity (Liu et al., 2021a). The combined model predicts which HLA molecule displayed a peptide that was observed to be immunogenic in a MIRA experiment, and uses machine learning predictions of peptide display for HLA alleles not observed or peptides not tested in MIRA data. Thus, all eight MHC class I peptides in our vaccine were previously observed to be immunogenic in data from convalescent COVID-19 patients (Liu et al., 2021a; Snyder et al., 2020). We further validated that all peptides are predicted to bind HLA-A*02:01 with high (≤ 50 nM) affinity using the NetMHCpan-4.1 (Reynisson et al., 2020) and MHCflurry 2.0 (O’Donnell et al., 2020) machine learning models. For inclusion in the assembled construct, the eight vaccine peptides were randomly shuffled, and alternate peptides were flanked with five additional amino acids at each terminus as originally flanked in the SARS-CoV-2 proteome.

We selected two additional peptides from the MIRA only vaccine design as negative controls that were not predicted to be immunogenic in HLA-A*02:01 transgenic mice, including by endogenous C57BL/6 mouse model MHC alleles (H-2-Kb and H-2-Db), by the MIRA/ML combined model, NetMHCpan-4.1, or MHCflurry 2.0.

#### MHC Class II

Three peptides from SARS-CoV-2 were optimized for predicted binding to H-2-IAb. We scored all SARS-CoV-2 peptides of length 13–25 using the sliding window approach of Liu et al. (2020) and a machine learning ensemble that outputs the mean predicted binding affinity (IC50) of NetMHCIIpan-4.0 (Reynisson et al., 2020) and PUFFIN (Zeng and Gifford, 2019). We selected the top three peptides by predicted binding affinity using a greedy selection strategy with a minimum edit distance constraint of 5 (Liu et al., 2021a) between peptides to avoid selecting overlapping windows. All three peptides were flanked with an additional five amino acids per terminus from the SARS-CoV-2 proteome.

#### Suitability for Protection Against Novel Variants

Table 1 shows the start and end amino acid positions of each peptide in its origin gene, including flanking residues for flanked vaccine peptides. For SARS-CoV-2, peptides are aligned to reference proteins in UniProt (Consortium, 2019) (UniProt IDs: P0DTC2 (S), P0DTC3 (ORF3a), P0DTC5 (M), P0DTC9 (N), P0DTD1 (ORF1ab)). We note that none of the SARS-CoV-2 peptides included in this vaccine were found to be mutated in the B.1.1.7, B.1.351, or P.1 SARS-CoV-2 variants (Rambaut et al., 2020; Davies et al., 2021; Tegally et al., 2021; Faria et al., 2021).

**Table 1.**
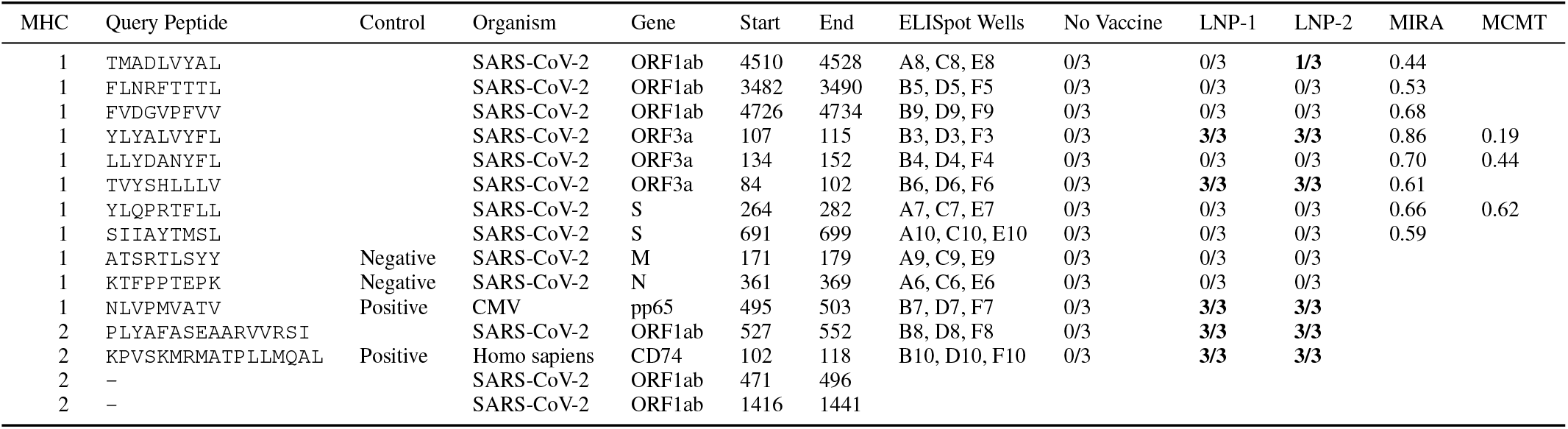
Immunogenicity of vaccine peptides. MHC class, ELISpot query peptide, control, peptide source, ELISpot wells, fraction of mice with positive ELISpot results out of 3, fraction of convalescent patients immunogenic for peptide (MIRA and MCMT). mRNA-LNP vaccines were formulated with either LNP-1 (ALC-0315) or LNP-2 (Tufts-113). Start and end values indicate peptide positions in the specified gene, including flanking residues if the peptide was flanked in the vaccine construct (see Methods). Two MHC class II vaccine peptides were not tested for immunogenicity during ELISpot. MIRA and MCMT are the fraction of convalescent SARS-CoV-2 patients that had an immunogenic response to each peptide in either a Multiplex Identification of T-cell Receptor Antigen Specificity (MIRA) assay (Snyder et al., 2020) or by mass cytometry–based multiplexed tetramer staining (MCMT) (Kared et al., 2021).

### 2.2 Construct design

Vaccine peptides were joined into a single polypeptide construct for mRNA-LNP delivery. Peptides were prepended with a secretion signal sequence at the N-terminus and followed by an MHC class I trafficking signal (MITD) at the C-terminus (Kreiter et al., 2008; Sahin et al., 2017), and joined by non-immunogenic glycine/serine linkers from Sahin et al. (2017). The construct also included positive control peptides previously shown to be immunogenic in HLA-A*02:01 and H-2-IAb mouse models (CMV pp65: NLVPMVATV for HLA-A*02:01 and Human CD74: KPVSKMRMATPLLMQAL for H-2-IAb). The assembled vaccine construct for our mouse experiments is shown in Figure 1.

**Figure 1.**
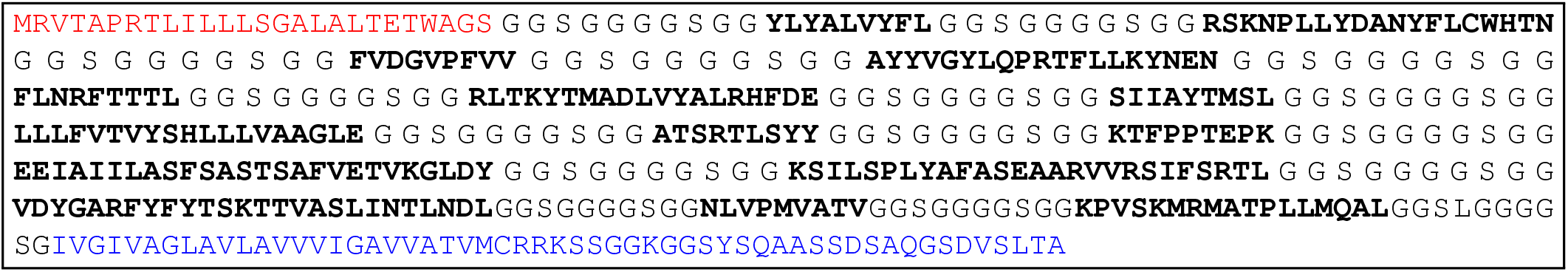
Assembled vaccine construct containing a secretion signal sequence (red), peptides (bold) joined by non-immunogenic glycine/serine linkers, and an MHC class I trafficking signal (blue).

Codon optimization for mouse expression of the vaccine construct was performed using the IDT Codon Optimization Tool (Integrated DNA Technologies), and the resulting nucleic acid sequence is provided in Figure S1.

### 2.3 mRNA-LNP vaccine formulation

RNA was synthesized by Trilink Biotechnologies as a modified mRNA transcript with full substitution of 5-Methoxy-U, capped (Cap 1) using CleanCap® AG and polyadenylated (120A). RNA containing lipid nanoparticles were prepared by dropwise adding an ethanol solution containing the mixture of active lipidoid, cholesterol (Chol), distearoylphosphatidylcholine (DSPC), and 1,2-dimyristoyl-rac-glycero-3-methoxypolyethylene glycol-2000 (DMG-PEG) at defined weight ratio to 25 mM sodium acetate solution. The Lipid : Cholesterol : DSPC : DMG-PEG weight rations were ALC-0315 16:7.44:3.35:1.86, Lipid Tufts-113 16:4.76:3:2.4. The mixed solution was then dialyzed using Thermo Scientific™ Slide-A-Lyzer™ MINI Dialysis Device (3.5K MWCO) to obtain the blank LNPs. The LNP/mRNA for vaccination was prepared by mixing 150 µg of blank LNP with 10 µg of mRNA.

### 2.4 Vaccine administration

A total of nine HLA-A*02:01 human transgenic mice (Jackson Laboratories Number 003475) were used. Three mice were unvaccinated controls, three mice were vaccinated with RNA formulated with the ALC-0315 lipid, and three mice were vaccinated with the same RNA formulated with the Tufts-113 lipid. The mice received their prime vaccination at Day 0 and a boost vaccination at Day 21. Each vaccination consisted of 100 µL of mRNA-LNP containing 10 µg of RNA injected subcutaneously at the base of the tail. Mice were sacrificed 14 days after the boost vaccination, and spleens were collected for ELISpot assay.

### 2.5 ELISpot analysis

Splenocytes were isolated, and 5 × 10^5^ splenocytes were placed in each well of a 96-well plate. Query peptides were obtained from GenScript at ≥95% purity. Query peptides were added at a concentration of 1 µg/mL, and the cell peptide mixture was incubated for 16 hours. Detection of interferon gamma positive spots was performed with a Cellular Technologies Limited CTL S6 FluoroSpot reader, and analyzed with ImmunoSpot 5.1.36 software.

### 2.6 MIRA and MCMT data analysis

Selected peptides were tested by other studies for their immunogenicity in convalescent COVID-19 patients whose HLA type included HLA-A*02:01. The study by Snyder et al. (2020) included 80 HLA-A*02:01 convalescent COVID-19 patients and tested peptides individually or in small pools with the Multiplex Identification of T-cell Receptor Antigen Specificity (MIRA) assay. Query peptides were first filtered to only consider those with predicted HLA-A*02:01 binding affinity ≤25 nM. The MIRA fraction in Table 1 is the number of individuals positive for a pool containing a query peptide divided by 80. The Kared et al. (2021) study evaluated 16 HLA-A*02:01 convalescent COVID-19 patient by mass cytometry–based multiplexed tetramer (MCMT) staining. The MCMT fraction in Table 1 is the number of individuals positive for a query peptide divided by 16.

## 3 Results

We tested the Figure 1 mRNA vaccine design and found that three vaccine SARS-CoV-2 peptides and the two control peptides were immunogenic (Table 1). The lack of normalization standards did not permit statistical analysis of the faction of T cells that were activated, or the difference, if any, of immunogenicity induced by mRNA-LNP 1 (ALC-0315) and mRNA-LNP 2 (Tufts-113). All positive control peptides were immunogenic. ELISpot wells corresponding to the data in Table 1 can be found for the control group, vaccine mRNA-LNP 1, and vaccine mRNA-LNP 2 in Figures S2, S3, and S4, respectively. The best ELISpot results from Group 2 (LNP-1, ALC-0315 formulated vaccine) and Group 3 (LNP-2, Tufts-113 formulated vaccine) are shown in Figure 2A and 2B, respectively.

**Figure 2.**
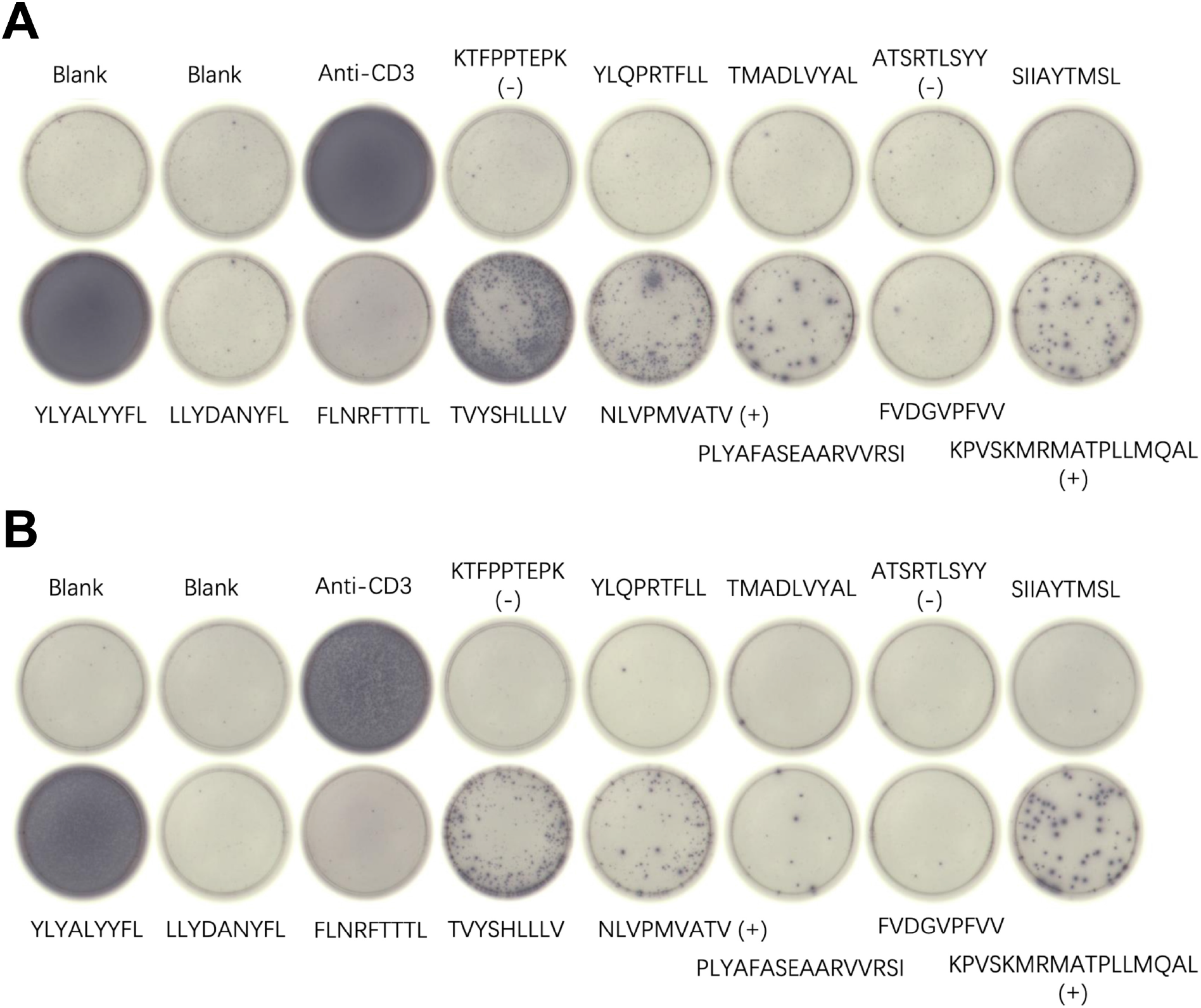
ELISpot results for indicated peptides resulting from an mRNA-LNP vaccine using (A) the ALC-0315 lipid (LNP-1, Mouse 2-3) and (B) the Tufts-113 lipid (LNP-2, Mouse 3-3). Blanks contained no added peptides, and positive controls (+) and negative controls (-) are indicated. Anti-CD3 antibody was added as a positive control to the indicated well.

## 4 Discussion

While we found two SARS-CoV-2 MHC class I peptides to be immunogenic, one highly so, five MHC class I peptides were not observed to be immunogenic in this study despite being immunogenic in certain HLA-A*02:01 convalescent COVID-19 patients. The lack of immunogenicity for these five peptides could be possibly a result of (1) incomplete processing of vaccine peptides in mouse HLA transgenics, as noted for other nucleic acid vaccines (Street et al., 2002), (2) the very high immunogenicity of YLYALVYFL may have competed against the immunogenicity of other peptides, (3) limitations in the T cell repertoire in the inbred HLA-A*02:01 mouse strains, and (4) the vaccine construct we created is too long for the vaccine to be optimally effective. However, we note that peptides both at the beginning and end of the construct were found to be immunogenic.

The observed immunogenicity of SARS-CoV-2 peptides YLYALVYFL and TVYSHLLLV (MHC class I), as well as PLYAFASEAARVVRSI (MHC class II) suggests that the vaccine may provide protection against high titers of SARS-CoV-2 viral infection.

## Acknowledgements

This work was supported in part by Schmidt Futures.

## Supplementary Information

**Figure S1.**
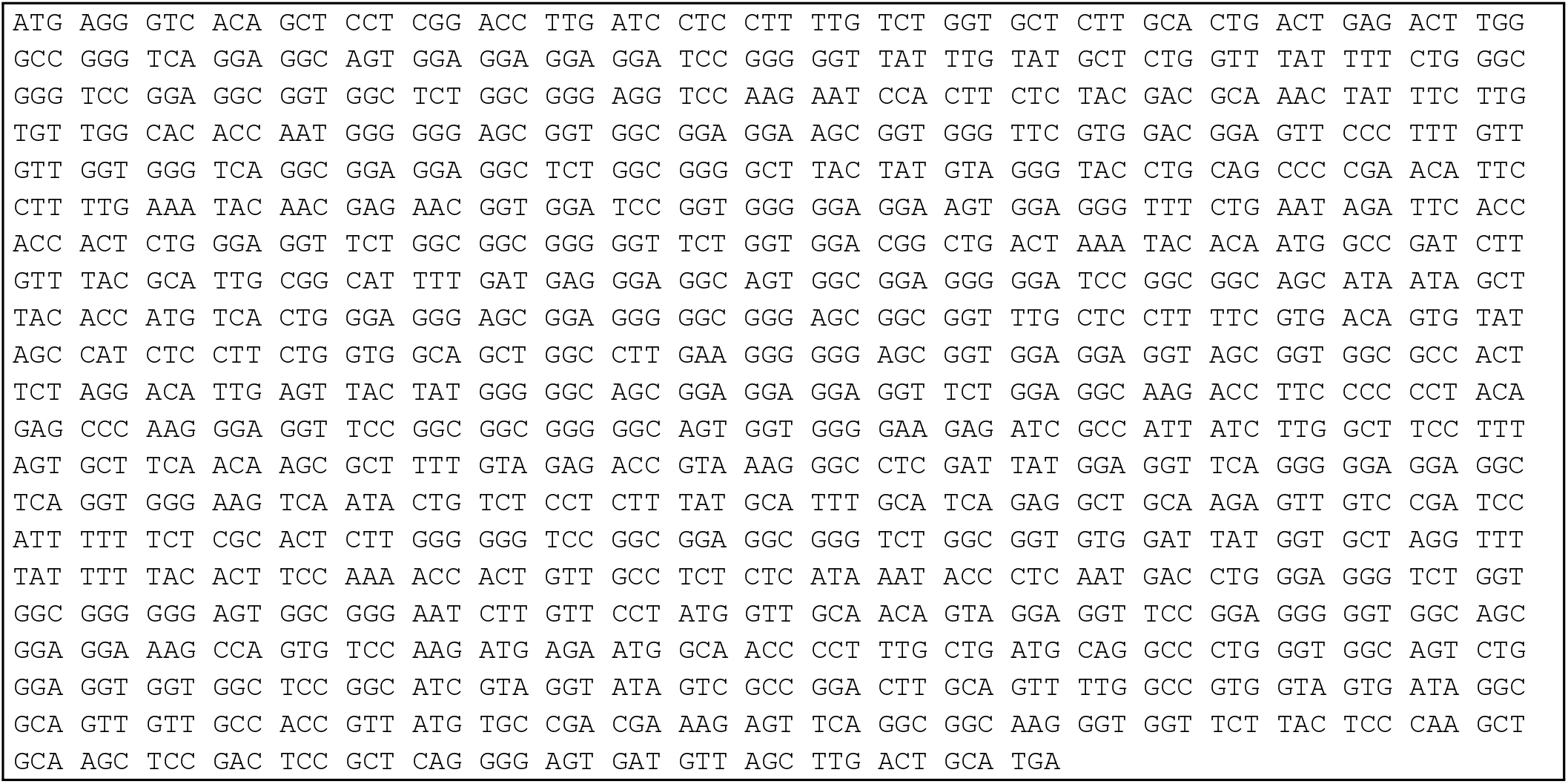
Nucleic acid sequence for assembled vaccine construct in Figure 1. Codon optimization for mouse expression was performed using the IDT Codon Optimization Tool (Integrated DNA Technologies).

**Figure S2.**
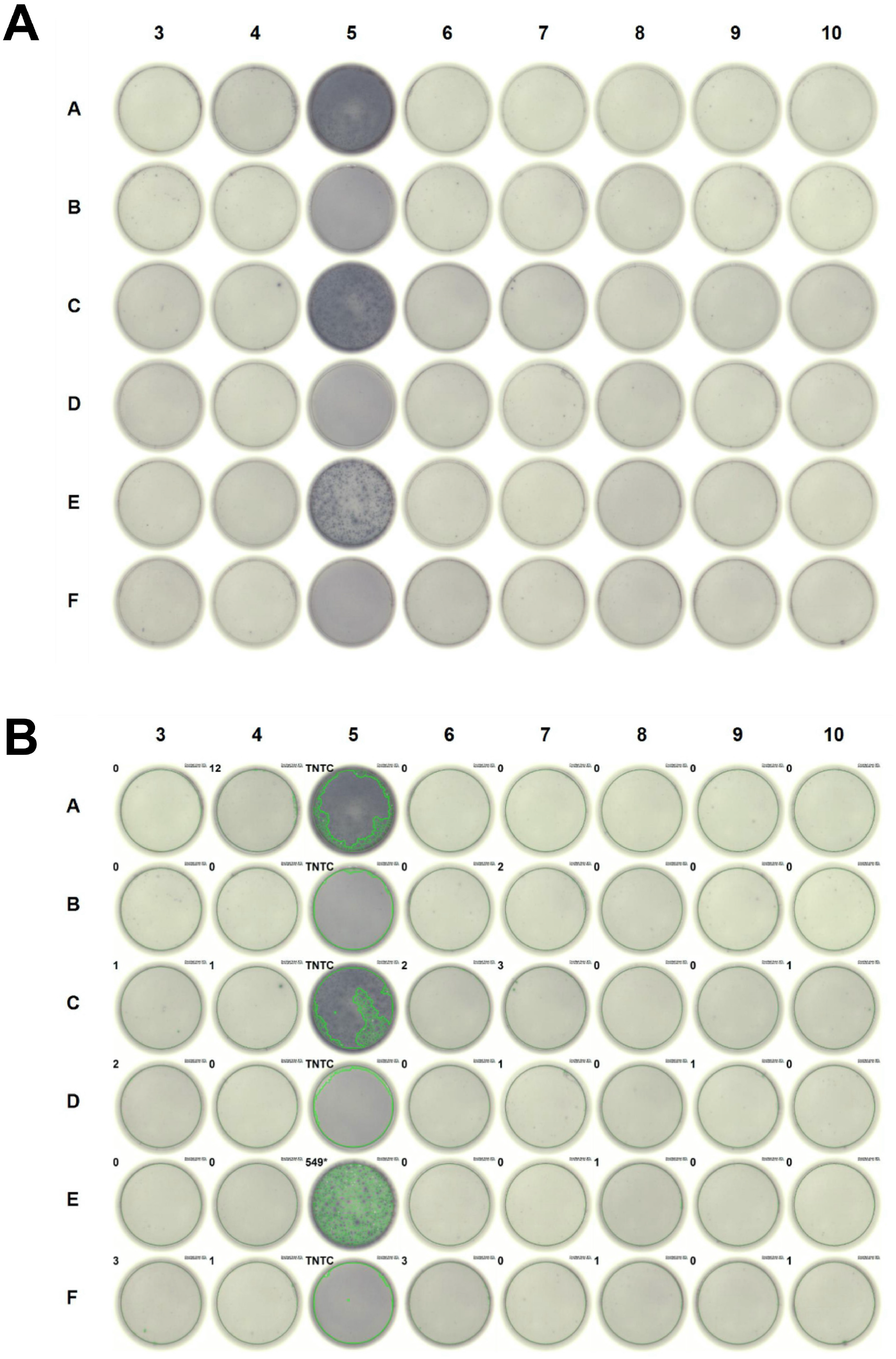
ELISpot plates for group: **No Vaccine**.

**Figure S3.**
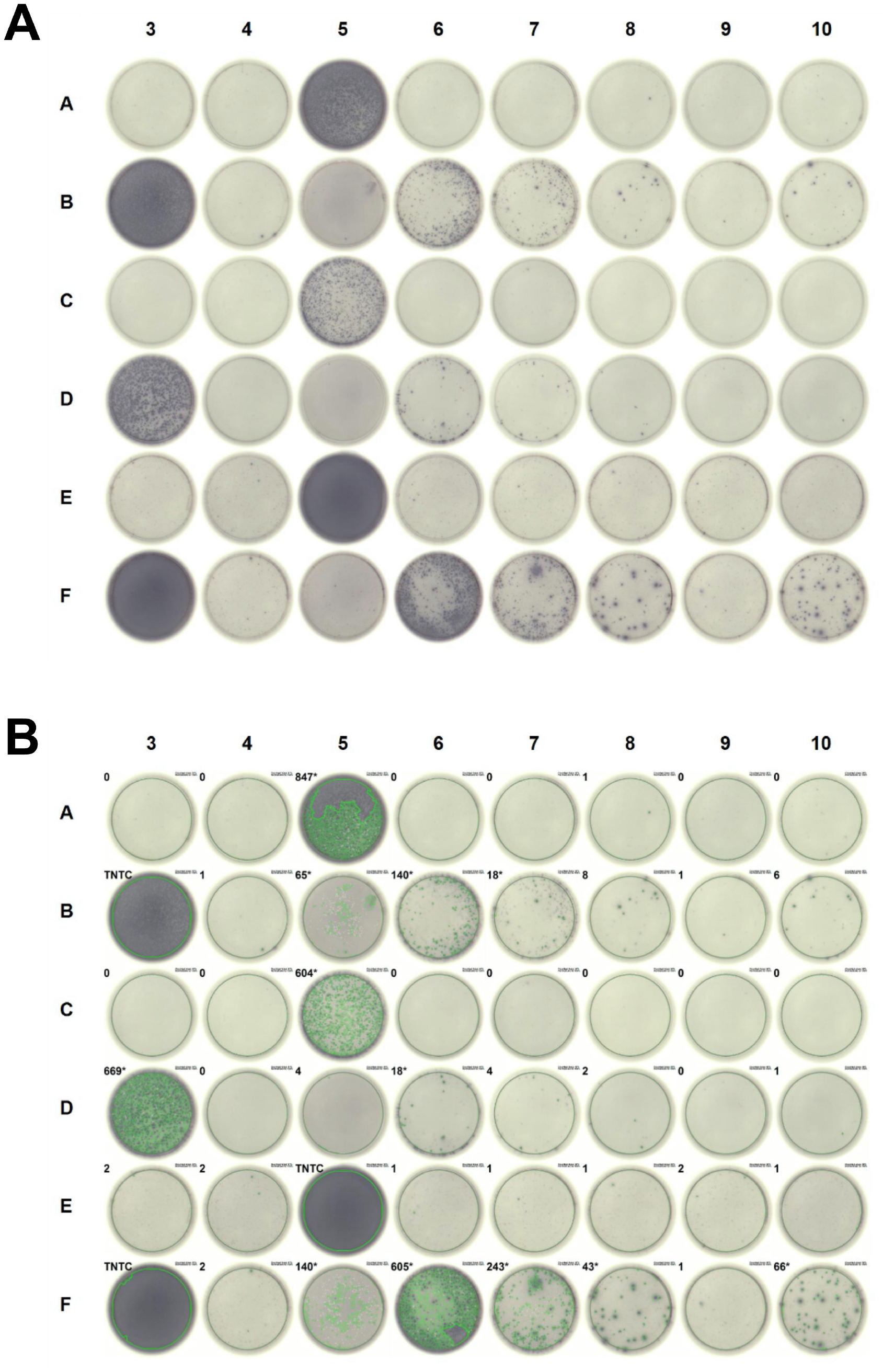
ELISpot plates for group: **LNP-1** (ALC-0315).

**Figure S4.**
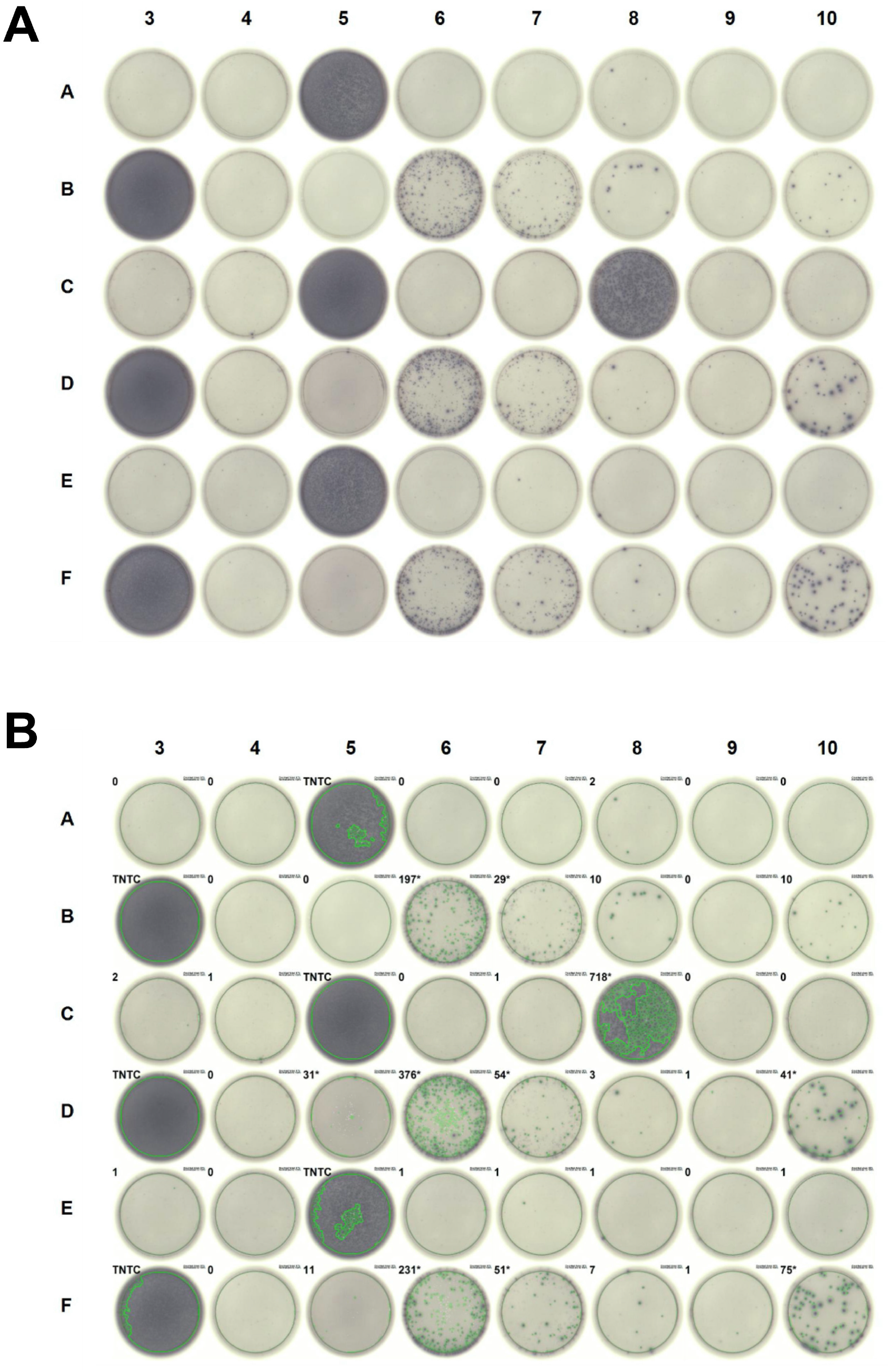
ELISpot plates for group: **LNP-2** (Tufts-113).

